# Gross anatomy of the gluteal and posterior thigh muscles in koalas based on their innervations

**DOI:** 10.1101/2021.12.13.472422

**Authors:** Sayaka Tojima, Hidaka Anetai, Kaito Koike, Saori Anetai, Kounosuke Tokita, Chris Leigh, Jaliya Kumaratilake

**Author notes:** **Corresponding author:** Sayaka Tojima. **Author contributions** Dr. Sayaka Tojima performed most tasks in this study (such as collaborative research with University of Adelaide, four koala dissections, and manuscript authorship and editing). Mr. Hidaka Anetai, Mr. Kaito Koike, Ms. Saori Midorikawa and Dr. Kounosuke Tokita performed one koala dissection and composed all figures. Mr. Chris Leigh provided important koala specimens and Dr. Jaliya Kumaratilake assisted in the dissection of koalas in Australia and contributed to the Discussion.

## Abstract

There are not many descriptions of the muscle morphology of marsupials, despite the fact that they should show diversity according to the adaptation and dispersal to a variety of environments. Most of the previous studies regarding the gross anatomy of marsupials were conducted in the 1800 - 1900’s, and many issues still remain that need to be reexamined. For instance, the muscle identification had been performed based only on their attachments and thus, muscle descriptions are often inconsistent among the studies. These classic studies often do not include figures or photographs, so the discrepancies in the descriptions of the muscles could only be verified by performing the muscle identification again with a more reliable method. This problem can be solved by performing muscle identification by innervation. This method, which focuses on the ontogenic origin of the muscle as opposed to the attachment site, is prone to individual and interspecies variation and is a common technique in recent anatomical research. This technique is more reliable than previous methods and is suitable for comparison with other taxa (i.e., eutherians). In this study, we first conducted muscle identification based on innervation in the gluteal and posterior thighs of koalas in order to reorganize the anatomical knowledge of marsupials. This is because the gluteus and posterior thighs of koalas are the areas where previous studies have been particularly inconsistent. We dissected five individual koalas and clarified discrepancies in previous studies, as well as investigated the unique muscle morphology and their function in koalas. Specifically, the koala’s gluteal muscle group is suitable for abduction, while the posterior thigh muscles are particularly suitable for flexion. In the future, we will update the anatomical findings of marsupials in the same way to clarify the adaptive dissipation process of marsupials, as well as to contribute to the understanding of the evolutionary morphology of mammals.

## Introduction

Marsupials are a group of mammals that diverged from eutherians, a group which includes us humans, about 130 million years ago. Marsupials were once distributed throughout the world, but lost out to eutherians and are now found only in the American and Australian regions [1]. Although the extant habitat of marsupials is more limited than that of eutherians, which are now widely distributed throughout the world, marsupials have adapted and dispersed during their evolutionary process, acquiring a variety of morphology. The koala, for example, is a well-known marsupial species; however, there is a lack of knowledge about their musculature which enables it to koala-specific locomotion or posture-maintenance.

In general, the number of studies investigating gross anatomy in marsupials are fewer than in eutherians. Additionally, most of the studies are classical, as they were conducted in the 1800-1900’s. Several classical studies have previously described the myology of koalas around the turn of the twentieth century [2-5]. However, even basic information about the muscular morphology in the koala was not fully clarified. Macalister [2] dissected the entire body of a female koala, but not all muscles were observed, and information on their attachments was insufficient. Sonntag [3] compared the anatomy of the wombat (*Phascolomys mitchelli*), koala, and phalanger (*Phalanger orientalis*); however, detailed information on each species was limited. Similarly, not all the attachment sites were described in the study by Macalister [2]. Elftman [4] compared pelvic region muscles among multiple marsupial species, producing drawings of the hip and thigh muscles. However, information about the attachments in dissected samples was not provided. Only Young [5] has summarized the attachments of koala muscles thoroughly. Another problem with the previous studies is the inconsistency in the description of muscles. For instance, Macalister [2] and Young [5] described that *m. gluteus medius* (GM) was bilaminar, while Sonntag [3] reported that there were no laminations. Thus, several pieces of information regarding muscle attachments remains debatable and hence requires updating and reappraisal. In other cases, muscles with similar attachment sites have sometimes been given multiple names that differ between species, leading to confusion [5-7]. The hind limb muscles of marsupials, especially the gluteal regions where multiple muscles overlap in layers, have been considered difficult to compare to those of eutherian mammals [7]. These inconsistencies occurred because the previous studies identified muscles only by their attachment sites.

To overcome this problem, muscle identification based on innervation of the muscles may be helpful because the nerve-muscle interactions exist, and the innervation reflects the developmental derivation of the muscles. Since the 1960’s, developmental biology has acknowledged these important interactions between muscle derivations and innervation, and it is important to consider the innervation to muscles, as it is essential to muscle functions, for two reasons: nerve–muscle interaction during the developmental process and specificity between nerves and muscles. Developmental biological studies have shown that normal muscular development and differentiation are strongly dependent on neural activities [8-12]. It is known that there is specificity between the nerves and muscles. Once a certain nervous fiber has connected to a particular muscle, the muscles cannot be connected with other nerves [13]. Furthermore, several anatomic investigations evaluating the derivations of muscles on the basis of innervation patterns have been conducted in humans and several mammals [14-29]. Thus, this method, which focuses on the ontogenic origin of the muscle, is a common technique in recent anatomical research. It is more reliable than previous methods and is suitable for comparison with other taxa (i.e. eutherians). As for marsupial anatomy, Koizumi [29] focused on the innervations and successfully identified the serratus anterior and scalenus muscles in koalas and compared the findings with those of cats.

Thus, in this study, we first focused on the muscles in the gluteal and posterior thigh regions of koalas, which have been difficult to identify and whose descriptions have been inconsistent, and attempted to re-identify and describe them based on their innervation. This study aims to update the findings of previous studies on the description of muscles in the koala gluteal and posterior thigh, and to provide insights that will contribute to the foundation of functional anatomical studies of marsupials.

The knowledge of the muscle morphology and function of koalas would provide important insights not only into the adaptive dispersal of marsupials, but also insights into the broad scale evolution of mammalian morphology. Koalas are arboreal like many other marsupial species, but they show a crucial morphological difference in that they have lost a tail, which would normally serve as a balancer in arboreal life. This is a very interesting morphology from a functional anatomical point of view; there is a hypothesis that slow arboreal locomotion might be an evolutionary factor for the loss of tails in some eutherians, including hominoids [30, 31]. Additionally, a very recent kinematic study showed that koalas have converged on the same strategies as primates with respect to climbing [32]. Therefore, this paper also aims to describe the unique muscle morphology of the koala and to discuss its function.

## Material and methods

### Koala specimens

Five adult koala specimens (four males and one female) were used in this study (Table 1). All animals had been previously fixed and stored in 10% buffered formalin after having their thoracic and abdominal cavities dissected and digestive organs removed. They were housed at the Discipline of Medicine, the University of Adelaide, South Australia. The animals were originally collected frozen from Cleland Wildlife Park under the Permit number K23749-12 from the Department of Environment, Water and Natural Resources, Government of South Australia. These specimens had previously been used for teaching comparative anatomy at the University of Adelaide, but their hind limbs were intact.

**Table 1.** Koala specimens used in this study.

| No. | Sex | Side |
| --- | --- | --- |
| 1 | Male | Both sides |
| 2 | Female | Both sides |
| 3 | Male | Both sides |
| 4 | Male | Right side |
| 5 | Male | Right side |

### Dissection

In this study, muscles of the gluteal region and posterior thigh and the nerves innervating them were dissected in detail. These are the muscles involved in the movement of hip and knee joints; thus, we hypothesized that the muscular morphology in these regions is responsible for the locomotion and/or vertical tree-hugging posture of koalas.

## Results

We successfully identified five types of muscles in the gluteal region and four types of muscles in the posterior thigh, based on their innervations and attachments. In this section, we described each of the muscles exposed from the superficial to deep layers. There were no variations in the innervations and attachments by individual or sex.

All koalas dissected in this study possessed eight lumbar and three sacral vertebrae. The nerves innervating each muscle were tracked back to their roots and identified (Figs 1-4). The sacral plexus in koalas comprised the most caudal branch of the sixth lumbar nerve (L6) to the second sacral nerve (S2).

**Fig 1.**
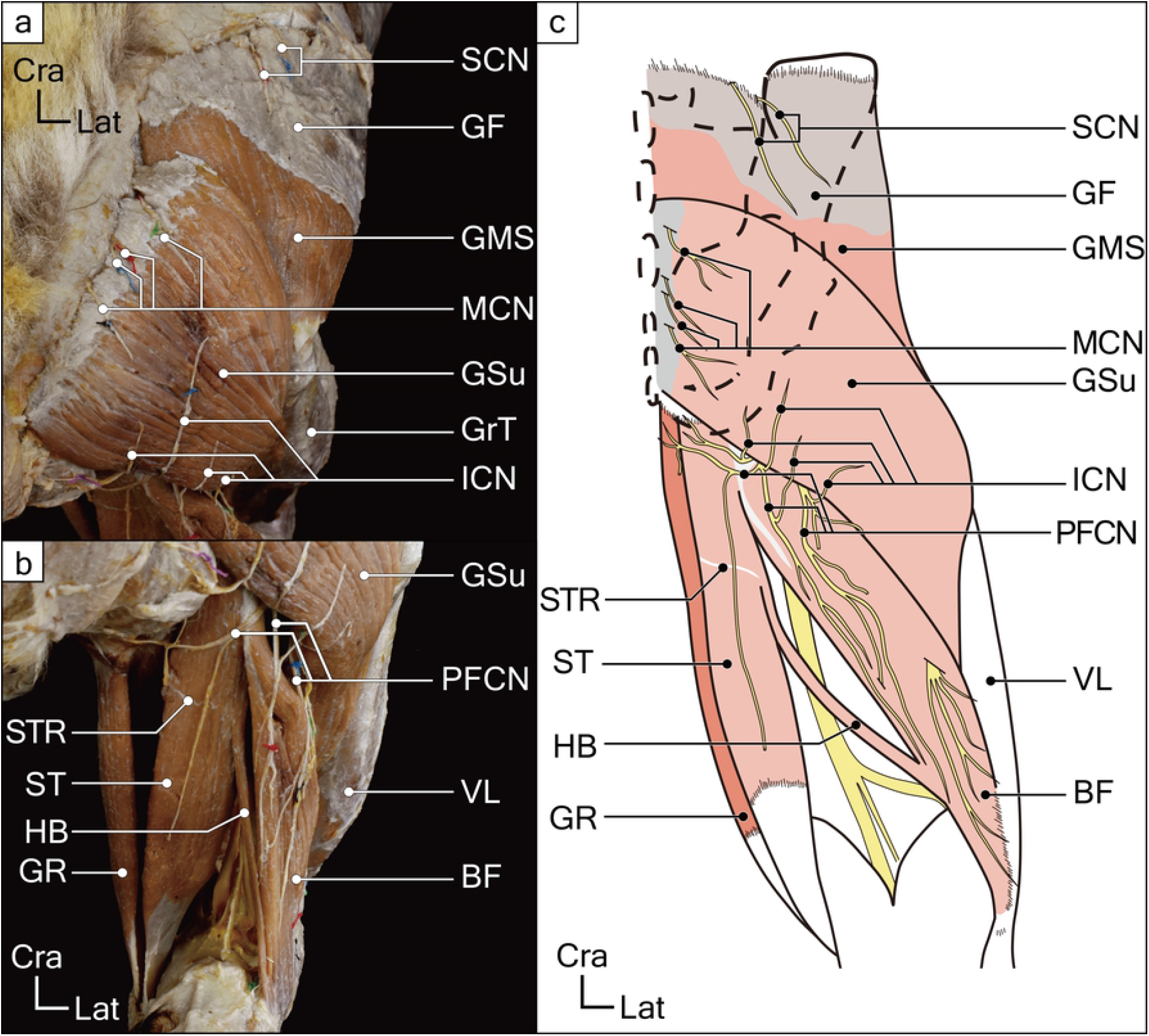
Superficial layer muscles of gluteal and posterior femoral regions in a koala right hind limb. (**a**) The superficial gluteal muscles, the gluteus superficialis (GSu) and superficial part of the gluteus medius (GMS), and some cutaneous nerves are shown. (**b**) The thigh flexors and some cutaneous nerves are shown. (**c**) Illustrated schematic drawing of the superficial layer muscles of gluteal and posterior femoral regions. Abbreviations: BF, biceps femoris; GF, gluteal fascia; GMS, superficial layer of the gluteus medius; GR, gracilis; GrT, greater trochanter; GSu, gluteus superficialis; HB, hamstring bundle; ICN, inferior cluneal nerve; MCN, middle cluneal nerve; PFCN, posterior femoral cutaneous nerve; SCN, superior cluneal nerve; ST, semitendinosus; STR, semitendinosus raphe; VL, vastus lateralis

**Fig 2.**
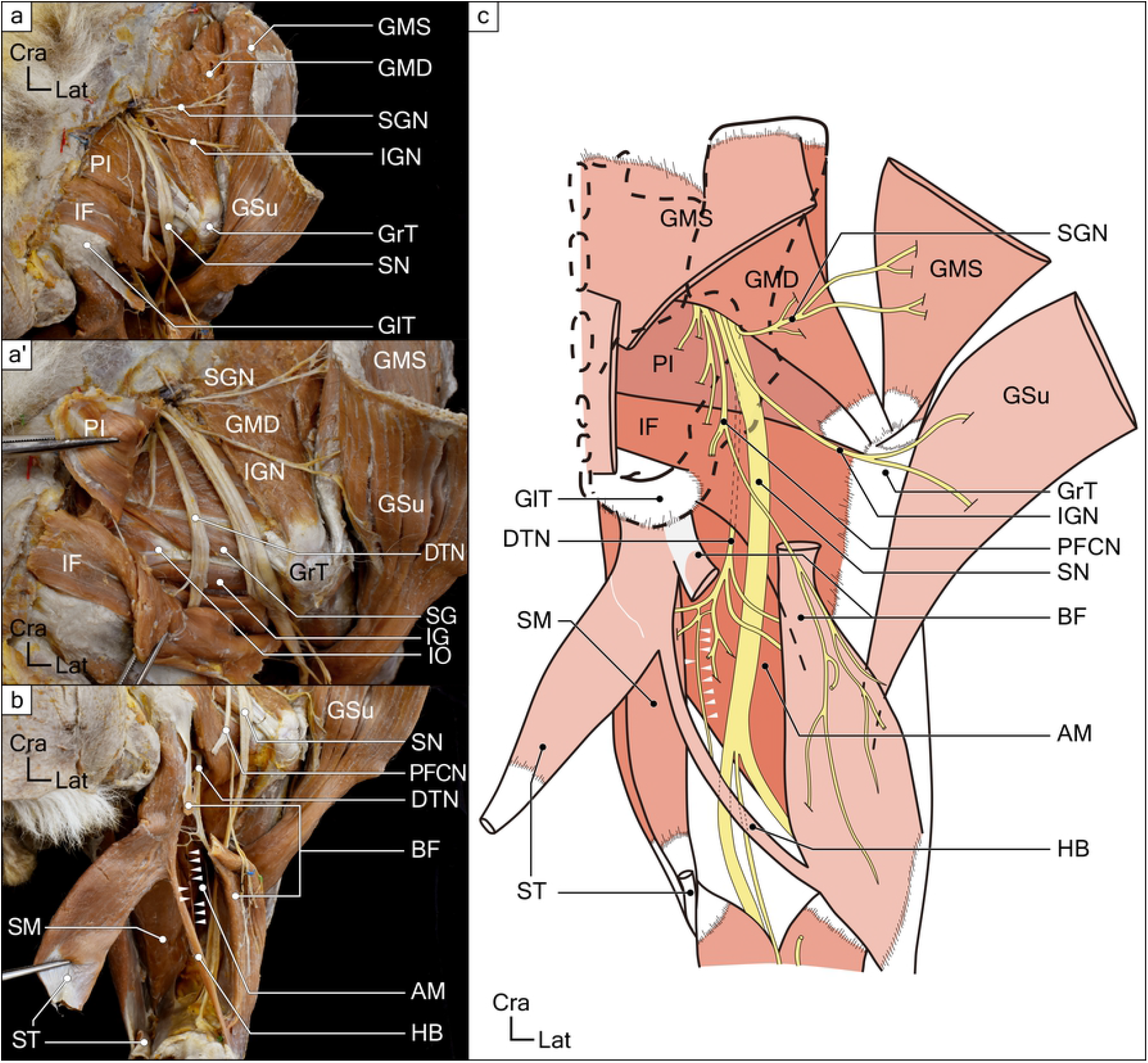
Deep layer muscles of gluteal and posterior femoral regions in a koala right hind-limb. (**a**) The deep gluteus muscles and several nerves including muscular branches are shown. (**a**) The deep tibial nerve (DTN) and deep gluteus muscles deep to the piriformis (PI) and ischiofemoralis (IF) are shown. (**b**) The innervating branches to the thigh flexors are shown. (**c**) Schematic drawing of deep gluteal muscles, thigh flexors, and innervating branches. White arrowheads indicate the innervating branch from the deep tibial nerve (DTN) to the hamstring bundle (HB). Abbreviations: AM, adductor magnus; BF, biceps femoris; DTN, deep tibial nerve; GlT, gluteal tuberosity; GMD, deep layer of the gluteus medius; GMS, superficial layer of the gluteus medius; GrT, greater trochanter; GSu, gluteus superficialis; HB, hamstring bundle; IF, ischiofemoralis; IG, inferior gemellus; IGN, inferior gluteal nerve; PFCN, posterior femoral cutaneous nerve; PI, piriformis; SG, superior gemellus; SGN, superior gluteal nerve; SM, semimembranosus; SN, sciatic nerve; ST, semitendinosus; VL, vastus lateralis

**Fig 3.**
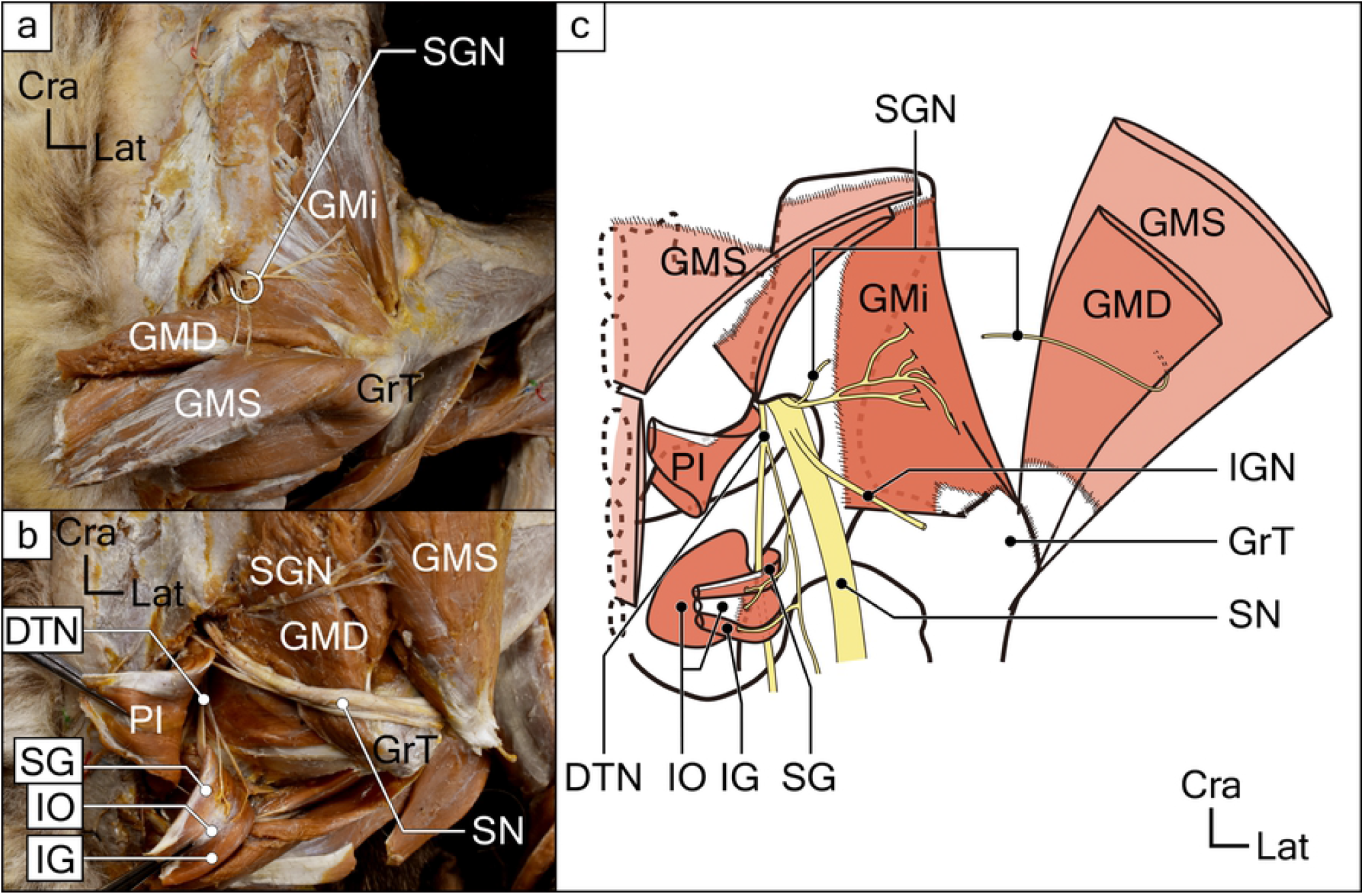
Deepest layer muscles of gluteus region and its innervations in koala right hip. (**a**) The superficial and deep layers of the gluteus medius (GMS and GMD), gluteus minimus (GMi), and the innervating branch from the superior gluteal nerve (SGN) are shown. (**b**) Gemellus and internal obturator muscles and supplying branch from the deep tibial nerve (DTN) are shown. (**c**) Schematic drawing of the deepest layer of gluteus muscles and its innervations. Abbreviations: DTN, deep tibial nerve; GlT, gluteal tuberosity; GMD, deep layer of the gluteus medius; GMi, gluteus minimus; GMS, superficial layer of the gluteus medius; GrT, greater trochanter; IG, inferior gemellus; IGN, inferior gluteal nerve; IO, internal obturator; PI, piriformis; SG, superior gemellus; SGN, superior gluteal nerve; SN, sciatic nerve

**Fig 4.**
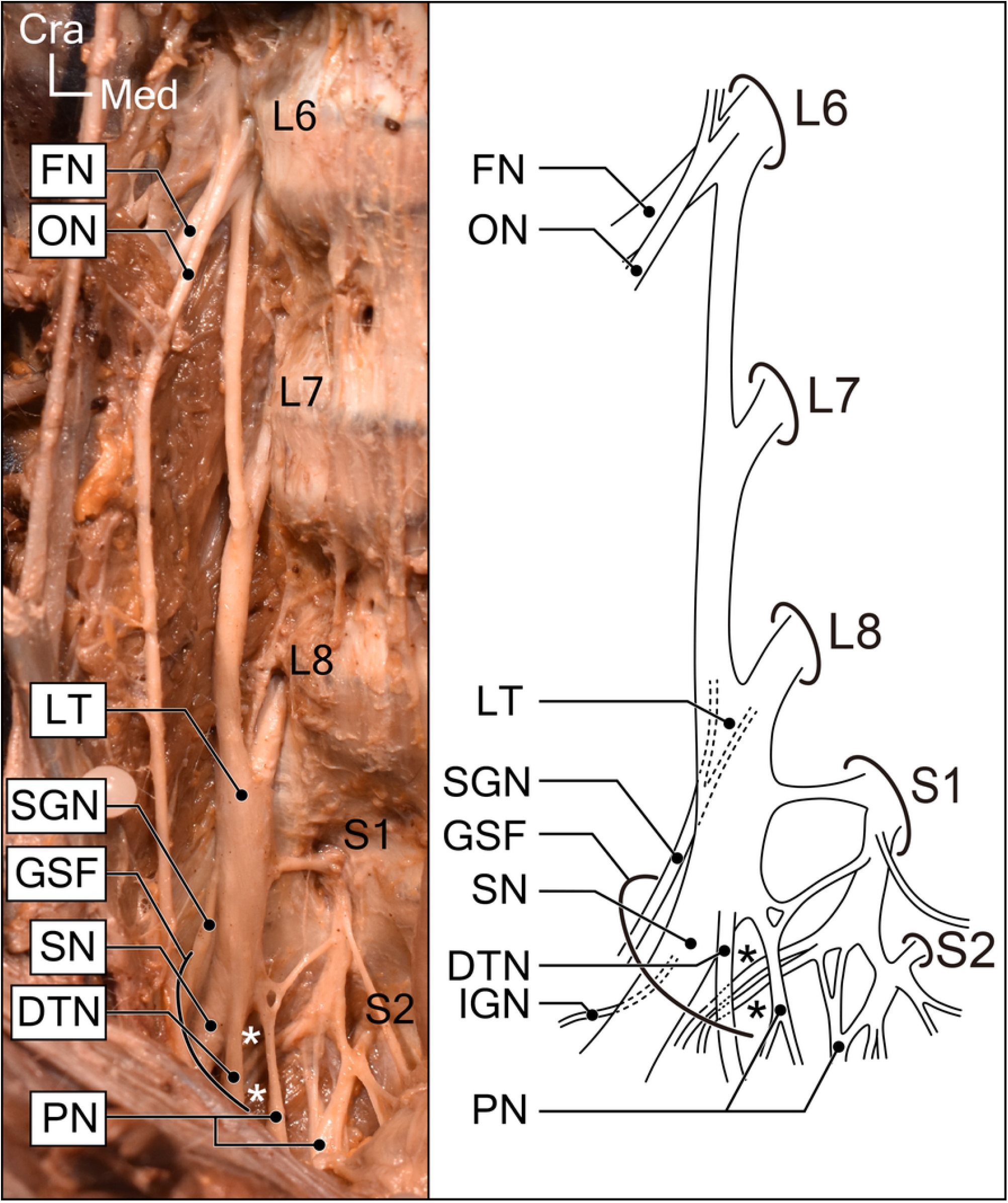
Right lumbosacral plexus in koalas. Photograph and illustrated corresponding schematic drawing. White and black asterisks indicate branches forming the posterior femoral cutaneous nerve and the innervating branch to the piriformis and ischiofemoralis after leaving the greater sciatic foramen. Abbreviations: DTN, deep tibial nerve; FN, femoral nerve; GSF, greater sciatic foramen; IGN, inferior gluteal nerve; LT, lumbosacral trunk; ON, obturator nerve; PN, pudendal nerve; SGN, superior gluteal nerve; SN, sciatic nerve

### Muscles in the gluteal region

#### 1. *M. gluteus superficialis* (GSu)

The GSu muscle was located in the most superficial layer beneath the superficial fascia. The muscle originated from the posterior aspect of the sacro-caudal region (from the third sacral vertebra to the third caudal vertebra) and was inserted onto the middle of the posterior surface of the femur shaft. The site of insertion extended approximately from 5 cm distal to the greater trochanter to 3 cm proximal to the distal end of the femur. There were no attachments to the ilium. These findings are consistent with the description of the gluteus maximus in several previous studies [2, 3, 5].

A branch from the sacral plexus (L8 to S1 nerves) entered the deep surface of the muscle. It was not a branch of the sciatic nerve, but possibly of the inferior gluteal nerve (Figs 1 and 2). Branches of the cutaneous nerves passed through the muscle near its origin (Fig 1).

#### 2. Superficial (GMS) and deep (GMD) M. gluteus medius (GM)

The GM muscle was located deeper than the GSu muscle. It originated from the posterior sides of the first to the third sacral vertebrae and the iliac crest and was inserted onto the lateral side of the greater trochanter. The muscular fascia separated the muscle into the GMS and GMD parts (Figs 2 and 3). The deep part originated along the shallow part of the posterior sacro-iliac ligament and was inserted onto the greater trochanter as a common tendon with the superficial component of the muscle.

The nerve that innervated this muscle originated from the L7 to L8 nerve roots, i.e., the superior gluteal nerve, entered the muscle from the deep surface and supplied both its GMD and GMS parts. Several branches of the cutaneous nerves supplying the skin on the superior part of the gluteal region passed through the cranial and medial parts of the GMS belly (Fig 1).

#### 3. *M. gluteus minimus* (GMi)

The GMi muscle, with a single belly, was located deeper than the GMD. It originated from the surface of the ilium and was inserted onto the greater trochanter. The nerve that supplied this muscle branched from the one that innervated the GM (the superior gluteal nerve). This nerve entered from the superficial aspect of the muscle belly.

#### 4. *M. piriformis* (PI)

The PI muscle was exposed after the transecting and reflecting of the GSu and GM muscles. It originated from the ventral side of the sacral body and was inserted onto the greater trochanter. The sciatic nerve (L6-S2) passed above this muscle, i.e., the suprapiriform foramen, and then descended (Figs 1-3). The nerve innervating this muscle branched from the sacral plexus (S1), close to the posterior femoral cutaneous nerve, and then entered the muscle belly from the superficial surface. No other muscle was observed to be homologous to the piriformis, except this muscle. Taking its attachments and innervations into account, it was identified as the piriformis muscle in koalas.

#### 5. *M. ischiofemoralis* (IF)

This muscle was located in the same layer as PI and was caudal to it. The origin was from the median sacral crest, sacrotuberous ligament, and the ischial tuberosity. The insertion was broad and expanded from the posterior edge of the greater trochanter to the shaft of the femur (i.e., just above the insertion of the GSu) (Fig 3).

The nerve innervating this muscle branched from the nerve supplying PI (S1), and then entered the muscle belly from the superficial surface at the superior proximal edge (Fig 3). This precluded finding a homologous muscle in other species. Furthermore, the nerves innervating *m. semitendinosus* (ST), *m. semimembranosus* (SM), *m. biceps femoris* (BF), and *m. adductor magnus* (AM) descended on the deep surface of this muscle, and we conveniently referred to it as the “deep tibial nerve (DTN).”

### Muscles at the posterior side of the thighs

Four muscles, which originated from the ischial tuberosity and were supplied by branches of the sacral plexus, were observed in the posterior aspect of the thighs. The three muscles formed a conjoint tendon and were inserted onto the antero-medial surface of the proximal tibia (Figs 1-3). The attachments and nerve innervations indicated that they were hamstring muscles.

#### 1. M. biceps femoris (BF)

The origin of this muscle was tendinous, from the ischial tuberosity. BF had a single head, without the short head found in its human counterpart. The muscle belly spread out like an isosceles triangle and became aponeurotic. This aponeurosis attached to the superior anterior aspect of tibia (i.e., from the proximal end to the tibial tuberosity) and continued to join with the conjoint tendon. Between this muscle and ST (described later), a muscle bundle connecting these two muscles was found and referred to as the “hamstring bundle” (HB) (Figs 2 and 3). Several branches of the posterior femoral cutaneous nerve descended onto the surface of this muscle. In addition, other cutaneous nerves passed through the muscle belly and innervated the disto-lateral area of the posterior thigh. The BF muscle was supplied by some branches of the DTN, as were the ST and SM (described later) (Fig 3)

#### 2. M. semitendinosus (ST)

This muscle originated from the ischial tuberosity, became tendinous at the distal part, and then joined the tendon of the gracilis muscle to form a part of the medial conjoint tendon. At the middle of the muscle belly, semitendinosus raphe (STr) was observed. A muscle bundle named the HB in this study, connecting the ST and BF, diverged from this muscle. Both the ST and HB were supplied by the DTN, branches of the sacral plexus. The supplying nerve entered the muscle from the deep surface.

#### 3. M. hamstring bundle (HB): a muscle bundle between ST and BF

This muscle connects ST and BF, and looks similar to the tensor fasciae suralis muscle in humans [23]. However, the directions of the muscle fibers are opposite; the fibers ran from ST to BF in koala (Figs 2 and 3), whereas those in the tensor fasciae suralis are reported to run from BF to ST. The nerve innervating HB was a branch of then DTN, which is a branch from the sciatic nerve, and this muscle is therefore a hamstring muscle. This muscle bundle was observed in six of the eight side sections of the koalas.

#### 4. *M. semimembranosus* (SM)

This muscle was located behind the ST muscle, and its origin was the ischial tuberosity. The tendon of this muscle was inserted onto the medial condyle of the tibia. Several nerve branches from the DTN entered the superficial surface of the middle part of the muscle belly (Fig 3).

## Discussion

This study provided two important results for understanding the gross anatomy of koalas. First, by focusing on the innervation as well as the attachment sites of the muscles, it became possible to identify the muscle better than in previous studies. In this way, we succeeded in organizing and updating the findings on muscle morphology in koalas, which had been inconsistent among previous studies. In addition, a muscle identification based on neuro-muscular specificity has enabled us to establish comparison with the eutherian, which has been considered difficult in the past. Among eutherian species, more numerous studies have been performed to clarify their gross anatomy. The comparison of muscle morphology and function with that of eutherians, which evolved by adapting to similar environments despite their distance in phylogeny, is very useful for considering adaptation and evolution in marsupials. This study is the first step in furthering the study of marsupials by enriching the anatomical knowledge that is the foundation of functional anatomy.

### Updated gross anatomy in the gluteal and posterior thigh regions of koalas

As stated above, the descriptions regarding several muscles were not consistent among previous studies. In addition, many of them were not accompanied by photographs or diagrams, so they could not be verified.

Inconsistencies in descriptions of muscles were particularly pronounced in the gluteal region, where multiple muscles overlap. While there was general agreement on the description of the GSu at the most superficial layer, there were differences in description at deeper levels: especially for the GM muscle, there was disagreement. As is mentioned in the introduction of this study, Macalister [2] and Young [5] have reported that GM of koalas were bilaminar, while Sonntag [3] had not observed the laminations. Our observations in this study supported the former observation, as we confirmed the layered structure of GM muscle in all six koalas. In addition to the koalas in this study, a case in which the GM is clearly divided into two layers has been reported in marsupial wolves [33].

As for the GMi, we observed it as an undivided single muscle. Similar to the description by Young [5], we found that the positional relationship between the PI muscle and the superior, inferior, and sciatic nerves was unique. Young [5] has described this muscle in koalas as “developed” and its attachments “as usual.” The attachment and innervation suggest that this muscle is the PI, but the three major nerves (superior and inferior gluteal nerves and the sciatic nerves) all emerged superior to the PI, i.e., the suprapiriform foramen, in the koala (Figs 2 and 3). In marsupials, there is only one previous study showing that the sciatic nerve exits from the suprapiriform foramen in echidna (*Tachyglossus aculeatus*) [34], and no similar case has been reported. The positional relationship between the PI muscle, superior and inferior gluteal nerves, and sciatic nerve in koalas is unique among mammals.

IF was also found to be inconsistent with the description of previous studies on koalas. In previous studies on marsupial anatomy, muscles with similar attachment sites are sometimes given multiple names that differ between species, leading to confusion (quadrutus femoris [5]; caudo-femoralis [6]; ischiofemoralis [7]). The attachment site and location of the IF observed in this study are consistent with those described by Young [5] as the quadratus femoris muscle, but the attachments and innervation are different from those of the quadratus femoris in eutherian mammals, including humans. First, the muscle observed in koalas is attached to both the sacrum and sacrotuberous ligament, and its origin is located more medially. The insertion was broader and located more distal to the trochanteric fossa. Additionally, the muscle of koalas had only a branch that entered from the posterior (superficial) surface of the muscle belly and did not have a branch that entered from the anterior (deep) surface as does the quadratus femoris muscle of many eutherian mammals, including humans. Finally, this muscle was located in a different layer (more superficial) than the superior and inferior gemellus and the obturator internus muscles. Considering the origin, insertion, innervation, and laminae, we concluded that this muscle is not identical to the so-called quadratus femoris. Based on the site of attachment, it is appropriate to call it the ischiofemoralis muscle.

As for the muscles of the posterior thigh, the types of muscles present and their mutual positional relationships were similar to those of previous studies and other eutherian mammals. The following two points are the exceptions. First, in a previous study, Sonntag [3] described that the BF and SM of koalas arise from a common tendon. Instead, we found that both muscles arise from ischial tuberosity, but from independent origins and not a common tendon (Fig 2). Next, we observed the presence of the HB (a muscle bundle connecting ST and BF) in six of the eight side sections of the koalas we observed. From the nerve innervation, it is certain that this is also part of the hamstrings, but since it is a thin muscle bundle, it does not seem to have any function. However, it has not been described in any previous studies.

Thus, in each of the gluteal and posterior thigh regions, we succeeded in clearing up several points of confusion from descriptions in previous studies. Moreover, the identification method based on innervation allows us to compare the function of the same type of muscle with that of eutherians, as described below.

### The function of the gluteal and posterior thigh muscles in koalas

The hind limb muscles of marsupials, especially the gluteal regions, have been considered difficult to compare to those of eutherian mammals [7]. However, the muscles observed in this study can be identified by their innervation, and their morphology and function can be compared with those of eutherian mammals. At present, there are few pieces of knowledge of muscle identification based on innervation in marsupials, making interspecies comparisons difficult. However, if new anatomical findings using this muscle identification method are accumulated in the future, it will be possible to make broad comparisons among marsupials, which will be useful in considering adaptive radiation in marsupials.

The results of this study suggested that the koala gluteal muscles may be adapted to the posture and locomotion of the koala hugging a tree with the hip abducted. First, in many eutherians, including humans, the GSu has an iliac origin, which allows the muscle to perform the functions of hip extension and external rotation. Although many marsupials also have an GSu origin in the iliac crest, this is completely lost in koalas. This suggests that the GSu in koalas acts mainly as a hip abductor and pulls the femur laterally and dorsally.

The fact that they have a two-layered GM muscle and an independent GMi also supports the adaptation of koalas to the tree-hugging posture and locomotion. According to previous studies, there are many cases in which the GM and GMi muscles are fused in marsupials. For example, the wombat (*Vombatus ursinus*) (considered to be relatively closely related to the koala), the Tasmanian devil (*Sarcophilus harrisii*) [2], and the marsupial mole (Notoryctidae, species unknown) have been reported to present fusion of the GM and GMi muscles [6], and Sonntag [3] noted that fusion of the muscles was observed in some marsupial species (species unknown). However, our study showed that koalas have a clearly two-layered GM and independent GMi. In particular, the GMS, which runs mediosuperiorly to lateroanteriorly on the dorsal side of the hip joint and inserts at the lateral side of the greater trochanter to cover the greater trochanter, is assumed to have an abduction function to pull the femur laterally and dorsally. In addition, the caudal part of the GMi (around two-thirds of the total muscle), which runs medially to laterally on the dorsal side of the hip joint and inserts at the great trochanter at the lateral side of the hip joint, is also likely to have an abduction function.

This is thought to be because these two types of muscles each perform important functions. The fusion of these muscles were reported only in terrestrial marsupial species. Perhaps, in terrestrial species, having one fused large muscle with a small range of motion may be more adaptive to the powerful back-and-forth movement of the hind limbs, which enhances propulsion. In contrast, arboreal marsupials, such as the koala in this study, may have been able to ascend and descend tree trunks of various thicknesses by increasing their range of motion through hip abduction and extension. These observations support the inference that each muscle exists independently.

Another major function of koala gluteal muscles is hip extension. The lower part of the GSu is expected to have an extensor function due to its similarity with the IF, described below, in terms of the running of muscle fibers. The GMD passes through the head of the hip joint and inserts at the great trochanter, indicating that it has an extensor function to pull the great trochanter towards the caudal side. In contrast, the cranial side of the GMi may play a role in flexion, which is antagonistic to this extension function; the cranial side of the GMi (approximately one-third of the total muscle) passes through the cranial side of the hip joint and attaches from the base of the great trochanter to the proximal lateral side of the diaphysis, where it pulls the femur cranially and medially, possibly performing flexion and internal rotation functions.

The fact that IF muscle is present instead of *m. quadrutus femoris* is also characteristic of koalas. The quadratus femoris muscle in many mammals including humans acts as an external rotator muscle. However, the ischiofemoralis of koalas is expected to act as a hip extensor rather than an external rotator, considering its attachment site and relationship with the hip joint. Since the IF travels along the dorsal and caudal sides of the hip joint from origin to insertion and reaches the femur, it is likely to pull the femur in the dorsal and caudal directions (hip extension). In addition, the lower part of this muscle inserts much like the adductor magnus muscle. Therefore, it is conceivable that it may also have a partial adductor function to prevent over-abduction of the hip joint.

On the other hand, the posterior thigh muscle was found to act as a hamstring, just like the other eutherians. A unique feature of the koala is that the pes anserinus is not formed due to the absence of the patella, and the conjugated tendon inserts broadly at the tibia. This is most likely due to the semi-flexed attitude of koalas, as mentioned by Young [5].

Thus, the muscle morphology of the gluteal and posterior thigh regions of koalas may reflect their unique locomotion and posture maintenance (cuddling trees, semi-flexed knee). Future studies on other muscle groups will reveal muscle morphology that reflects the ecology of koalas, including upright posture in trees, which is different from that of other marsupials.

## Conclusions

In this study, we made important contributions to understanding gross anatomy in marsupials. Muscle identification based on innervation is a method that provides a stable standard for muscle identification across species. By using this identification method, we have established the basis for comparative anatomy between marsupials and eutherian mammals. Although straightforward comparisons between them are difficult due to unique marsupial adaptations and evolution, the use of muscle identification based on innervation allows the establishment of a clear correspondence of anatomical findings. This allows us to better infer muscle morphology and function, and depict various characteristic musculoskeletal morphologies. For example, this study found that the gluteal muscle group of the koala has excellent abduction and extension functions, and that the posterior thigh muscle group corresponds to the koala’s knee-bending posture. However, this study covered only a small part of koala muscles. In the future, we believe that by taking systemic anatomical findings and comparing with those of eutherian mammals, we will be able to clarify the unique anatomy of the koala more comprehensively.

## Acknowledgements

We sincerely thank Dr. Masatake Kai for his constructive suggestions and English corrections in improving this paper.

